# Single-cell sequencing reveals genome streamlining and functional diversity in ecologically dominant marine protists

**DOI:** 10.64898/2026.04.17.719054

**Authors:** David López-Escardó, Aleix Obiol, Guillem Marimon, Xabier López-Alforja, Dolors Vaqué, Irene Forn, Ramiro Logares, Sheree Yau, Òscar Fornas, Manuel Martínez-García, Ramon Massana

## Abstract

A large fraction of marine protists, particularly the smallest ones, belong to uncultured lineages that lack genomic data, limiting insights into their ecological roles and evolutionary strategies. Here, we generated 325 single-cell amplified genomes (SAGs) from 2-5 µm planktonic protists, which resulted in 147 genomes from dominant marine species at varying levels of completeness (40 of them >50%). These genomes matched the *in situ* community, with Prymnesiophyceae, Mamiellophyceae and Chrysophyceae dominating pigmented cells and MAST, Choanoflagellata and Picozoa prevailing among heterotrophic colourless cells. This resource allowed us to describe the genomic architecture of marine protist species, and revealed a pronounced genome streamlining in ecologically successful lineages. Comparative analyses highlighted unique functions enriched in photosynthetic and heterotrophic taxa (including motility, signal transduction, digestion and secondary metabolism), and revealed a broad distribution of gene families with adaptive traits such as polyketide synthases and rhodopsins. This large-scale single-cell genomics dataset provides a mechanistic foundation for investigating functional diversity, ecological strategies and genome evolution in the ocean.

## Background

Protists - unicellular eukaryotes - are extraordinarily diverse in terms of taxonomy, morphology and function, and their biomass in marine ecosystems rivals that of animals and prokaryotes^1^. Photosynthetic protists drive primary production, while heterotrophic forms act as predators, parasites and decomposers that regulate microbial populations^2^. Collectively, protists shape global biogeochemical cycles and channel inorganic and organic carbon to upper trophic levels or into the deep ocean^3^. Despite their ecological relevance, the full picture of marine protist diversity remains incomplete, largely because key lineages resist cultivation^4^. In the last decades, culture-independent molecular surveys unveiled this hidden diversity, which was especially critical for heterotrophic protists. Seminal studies based on 18S rRNA genes uncovered novel lineages such as Marine Stramenopiles (MAST)^5^, predators of prokaryotes; Marine Alveolates (MALV)^6^, parasites of dinoflagellates, radiolarians, and animals; and Picozoa^7^, whose trophic role still remains unclear. The large uncultured component of protist diversity results in a major genomic gap that hinders the mechanistic interpretation of ecological functions and evolutionary trajectories^8^, and limits predictions of how these species may respond to a changing ocean^9^.

Metagenome-assembled genomes (MAGs) have been proposed as a solution to the microbial genome gap, and large numbers of high-quality prokaryotic MAGs have been generated^10^. Similar efforts for microbial eukaryotes lag behind, and even with massive datasets like those generated by the TARA Oceans expedition, eukaryotic MAGs are often incomplete and fail to recover dominant taxa^11^. This suggests that MAG reconstruction is more challenging for eukaryotes than for prokaryotes, given their larger genomes, distributed across multiple chromosomes, and rich in repetitive regions. In this context, single-cell genomics (SCG) offers a promising alternative for accessing the genomes of uncultured protists. The approach for small protists resembles that used for prokaryotes^12^: cells are isolated by fluorescence activated cell sorting (FACS), genomic DNA is amplified, and the resulting material is sequenced and assembled into single-cell amplified genomes (SAGs). SCG typically yields fragmented and partial protist genomes, but completeness can be improved by coassembling sequences from conspecific cells^13^. SCG has already been applied to place uncultured eukaryotic species into multigene phylogenies^14,15^, to study interactions within single cells^16,17^, to recover mitochondrial genomes^18^, and to characterize the gene repertoires of groups such as MASTs^19^, choanoflagellates^20^, and ciliates^21^. However, the SCG pipeline has rarely been applied to target of the entire protist community and still requires improvements with respect to genome completeness and integrity.

Here we applied SCG to expand the genomic data for underexplored yet abundant marine protists. We targeted small planktonic cells (2-5 µm) from the Blanes Bay Microbial Observatory (BBMO), a northwestern Mediterranean station with long-term molecular records^22^. We sorted live cells to avoid potential damage during cryopreservation, and sequenced pigmented and colourless cells of different sizes. The 325 sequenced single cells yielded 147 genomes, as several cells were from the same species and were coassembled. These genomes represent previously unsampled ecological players, provide a detailed picture of the *in situ* protist diversity, and recover a substantial fraction of BBMO metagenomic reads. This dataset further allowed us to explore genome architecture across completely uncharacterized evolutionary lineages, including evidence for extensive streamlining^23^, identification of functions enriched in distinct trophic guilds, and analysis of the distribution of key gene families. Overall, our SCG collection constitutes a significant step toward resolving the ecological roles and evolutionary patterns of marine protists.

## Results

### A robust single-cell genomics approach for the smallest marine protists

Marine protists distribute along a logarithmic size spectrum^2^, with cells <5 µm being highly abundant and poorly characterized^4^. We established a simple and reliable SCG workflow capable of capturing the full diversity of these small protists, one cell at a time. In a first sampling (May 2018), we optimized the flow-cytometric identification of protist cells in a freshly collected seawater sample based on chlorophyll autofluorescence for phototrophs (Fig. 1a) and DNA staining for heterotrophs (Fig. 1b). Within each group, we sorted different size fractions, pico-phototrophic protists (pico-PP) and two populations of heterotrophic protists (pico-HP and nano-HP). Heterotrophic cells later identified as belonging to the same species exhibited very similar cytometric signatures (Fig. 1c), indicating that specific cell and genome sizes yield reproducible phenotypic fingerprints that could enable targeted, cytometric-based isolation strategies in future work. For whole genome amplification (WGA) from single cells we employed an optimized MDA-based protocol^24^, which is effective but produces partial amplifications^13^. WGA was positive for 75% of the sorted cells (60% in pico-PP, 78% in pico-HP and 83% nano-HP), a success substantially higher than in earlier work using cryopreserved cells (27%)^25^. This improvement is likely due to the use of fresh samples, which is only possible when cytometry facilities are near the sampling site. We next evaluated the utility of PCR screening prior to sequencing. Direct PCR with 18S rDNA primers succeeded in 50% of cells, with particularly low recovery in pico-PP (Fig. 1d). In contrast, Illumina assemblies yielded the 18S rDNA for nearly all PCR-positive cells and for cells in which PCR had failed (Fig. 1e). Many cells without detectable rDNA still produced usable genomic data (see later). Thus, 18S PCR screening can bias the selection of SAGs for sequencing and provides limited value when the aim is to describe the whole protistan community.

**Fig. 1.**
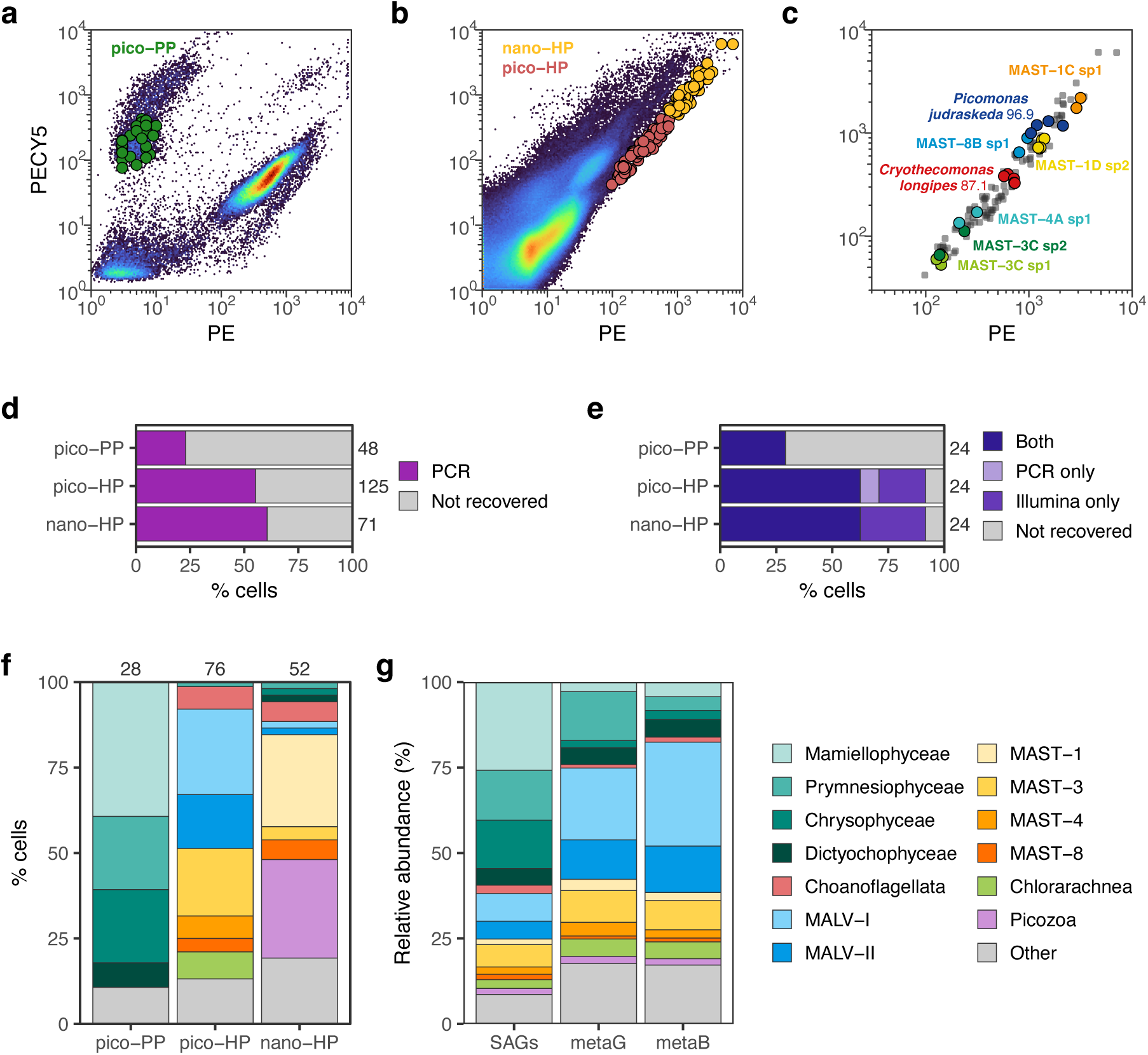
Performance of single-cell genomics for recovering marine pico- and nano-protists. Analyses were conducted on a BBMO sample (May 2018) sorted a few hours after collection. **a,** Flow cytogram of an unstained aliquot showing *Synechoccoccus* (right) and phototrophic protists (PP; left); large dots indicate sorted pico-PP cells. **b,** Flow cytogram of a DNA-stained aliquot highlighting colourless heterotrophic protists (HP) along the lower-right diagonal; large dots mark sorted pico-HP and nano-HP cells. **c,** Taxonomic identification of selected HP cells. **d,** PCR recovery of the 18S rDNA gene from cells with successful WGA. **e,** Recovery of the 18S rDNA gene in 24 sequenced cells per sort (also PCR-verified). **f,** Taxonomic affiliations of cells from the three sorted populations. **g,** *In situ* protist community composition inferred from a composite of SAGs across the three sorts and from metagenomic (metaG) and metabarcoding (metaB) datasets.

Taxonomic classification using 18S rDNA and phylogenomics revealed the expected diversity in each sorted fraction (Fig. 1f). Prymnesiophyceae, Mamiellophyceae, and Chrysophyceae dominated the pigmented cells, whereas colourless cells included mainly MAST, MALV, Choanoflagellata and Picozoa. HP fractions showed distinct taxa, with pico-HP enriched in MALV-I/II and MAST-3/4, and nano-HP enriched in MAST-1 and Picozoa. Comparison with metabarcoding and metagenomic datasets from the same sample showed that the normalized SAG-derived community composition closely matched *in situ* diversity (Fig. 1g), except for the overrepresentation of photosynthetic taxa, which is consistent with their low rDNA operon copy numbers and their typical underrepresentation in rDNA surveys^26^. Overall, direct Illumina sequencing of SAGs, without PCR pre-screening, proved to capture, without biases and accurately, the protist diversity of the sample. We therefore adopted this approach in a subsequent effort to expand the genomic catalogue of key microbial eukaryotes.

### New genomes of marine protists from single cells and coassemblies

Sequencing and assembling from BBMO single-cell sorting efforts in May 2018 and September 2020 yielded 325 SAGs of sufficient quality for downstream analyses (Table S1). Initial assemblies (contigs >1 kb) had an average size of 14.83 Mb, an N50 of 13.0 kb and 29.9% of completeness. Most contigs >3 kb derived from nuclear eukaryotic genomes (94.8% on average), with a few originating from prokaryotes (3.2%), organelles (0.6%) or unclassified sources (1.4%). SAGs belonging to the same species were identified through identical 18S rDNA and high average nucleotide identity (ANI), and their reads were then coassembled. Individual SAGs used in coassemblies exhibited very high ANI (98.1% on average) and genomic fragments peaked in the 99–100% bin for all but 3 cases (Fig. S1a). This supported conspecificity and validated the coassembling strategy. These plots contrasted with those of closely-related species, which typically exhibited a peak at 80-90% (Fig. S1b). The final dataset comprised 147 genomes (Table S2), 99 derived from single cells (SAGs) and 48 from coassembled cells (COSAGs), and their assembly features are shown in Fig. 2a. The final genomes averaged 23.3 Mb in size and 37.6% in completeness, and these values doubled in COSAGs (31.9 Mb and 57.6%) compared with SAGs (19.2 Mb and 27.9%). N50 values were also higher in COSAGs (16.6 kb) than in SAGs (10.9 kb). We predicted an average of 10,664 protein-coding genes per assembly.

**Fig. 2.**
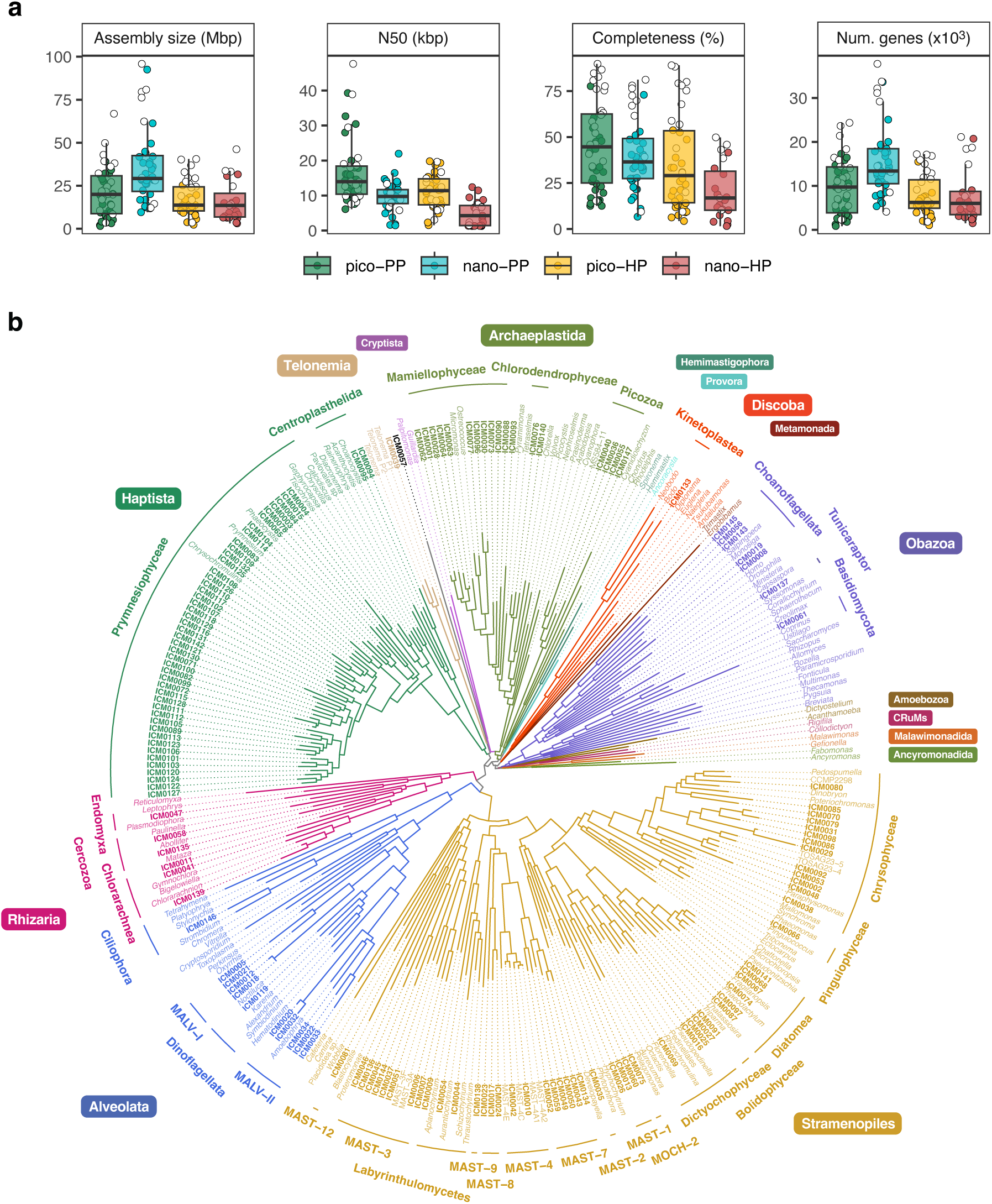
Assembly statistics and phylogenomics of the 147 protist genomes. **a**, Distributions of assembly size, N50, BUSCO completeness, and gene counts, separated by the sorted cell populations; white dots denote values from coassembled genomes. **b,** Multigene phylogeny of the new genomes and selected reference genomes, inferred with PhyloFisher using 240 conserved marker proteins. The full tree with all reference sequences and bootstrap values is shown in Fig. S2. Branches and species names are coloured by supergroup (indicated by coloured boxes), and detailed taxonomic groups with SAGs are displayed in the outer annotation ring.

Taxonomic profiling of the dataset highlighted both high diversity and uneven species richness across groups (Table 1). For example, there were four times as many species in Prymnesiophyceae as in Mamiellophyceae, despite the two groups having a similar number of cells. An even more striking contrast was seen between MALVs (9 cells and 9 species) and Chlorarachnea (38 cells and 3 species). A phylogenomic tree of our new genomes together with reference genomes recovered the eukaryotic tree of life, showing monophyly for the supergroups (except Archaeplastida) and for all lower-rank groups (Fig. 2b; Fig. S2). We obtained 20 genomes with more than 70% completeness (Table S2), including members of the environmental clades MAST-3 and -4, and novel lineages within Chrysophyceae, Choanoflagellata and Dictyochophyceae (<94% 18S rDNA similarity to its closest cultured relative). We also obtained the complete genome of *Minorisa minuta*, a species of particular ecological and evolutionary interest, as it is a colourless bacterivore at the base of a chloroplast-containing lineage^27^ and whose culture was lost prior to sequencing. With lower completeness, our dataset provides the first genomic data of several marine groups (only transcriptomic data existed for some of them), such as MALV-I, MAST-12, MOCH-2, Centroplasthelida, Dictyochophyceae, loricated and acanthoecida Choanoflagellata, and Telonemia. Only two species were not well assigned, ICM0093 as an unresolved Chlorophyta, and ICM0057 as an unknown lineage that branched deeply to Telonemia with low support. Thus, our dataset expands the genomic diversity of major marine eukaryotic groups, including novel groups that can be key to reconstruct eukaryotic evolution. In that sense, opisthokont SAGs could contribute to better reconstruct holozoan phylogeny and the origin of animals^28^.

**Table 1.**
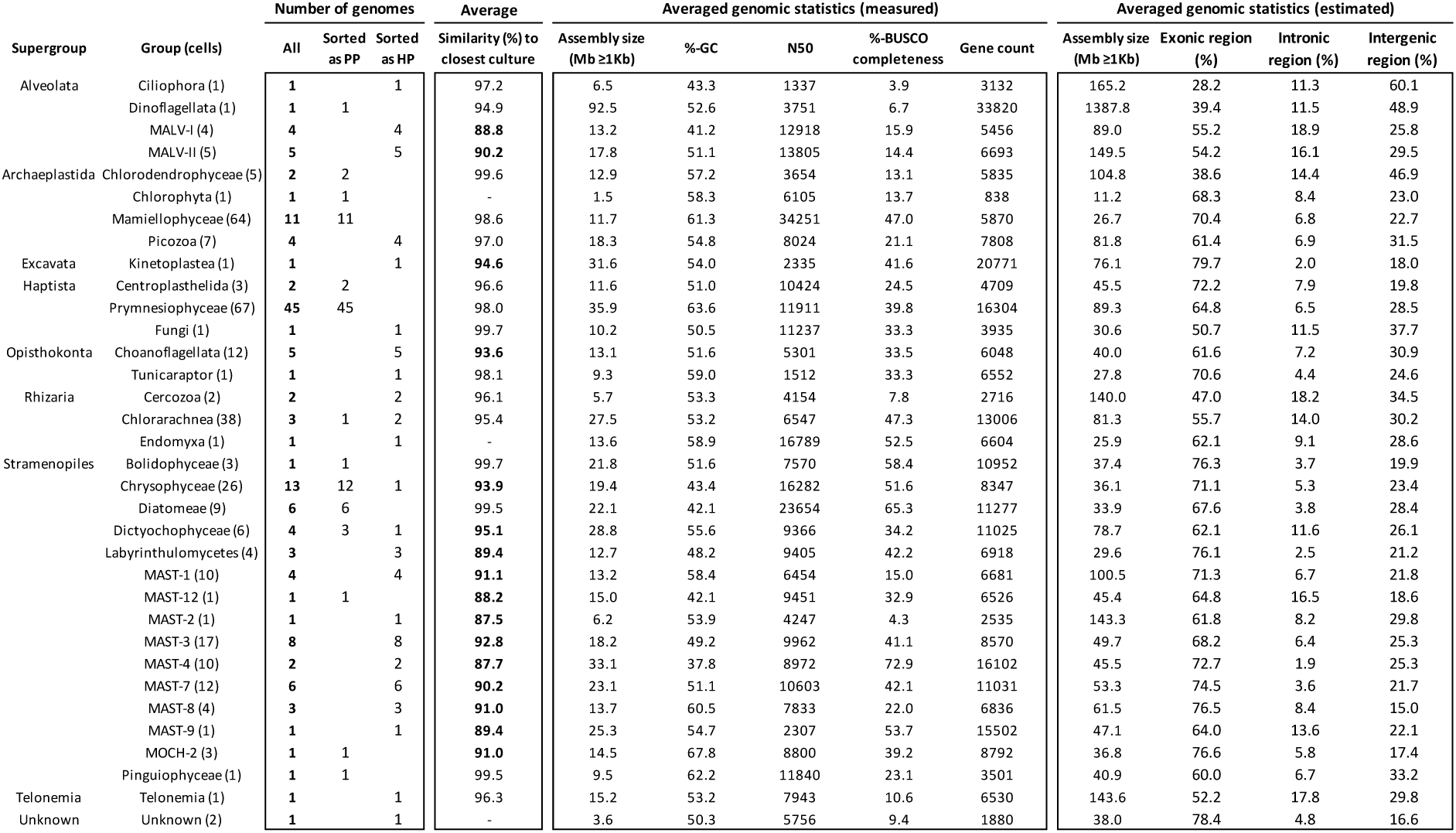
Overview of the 147 final genomes. For each taxonomic group (number of cells in parentheses), the table reports the number of genomes obtained from PP and HP sorts and the average 18S rDNA similarity to the closest cultured relative. Averaged measured features are shown, including assembly size, %-GC, N50, BUSCO completeness, and gene counts, together with the estimated genome size and the predictions of exonic, intronic and intergenic regions.

### SAGs account for dominant and widespread marine species

The ecological relevance of the species represented by our new genomes were evaluated using omics datasets. We first examined metabarcoding surveys targeting the V4 region of the 18S rDNA gene, including the BBMO time series, the coastal Ocean Sampling Day (OSD) dataset^29^, and the EukBank collection^30^. Searching for identical ASVs (Amplicon Sequence Variants) in the metabarcoding datasets was done in 109 genomes that had this region, and most found an identical match, 104 in BBMO, 93 in OSD and 92 in EukBank. These ASVs were abundant in the marine plankton, rare in marine sediments, and absent from freshwater and soils (Fig. S3), indicating that we retrieved the genomes of marine planktonic species. A detailed comparison was done in two datasets targeting small marine protists (Fig. 3a). ASVs corresponding to our genomes were common in the two datasets, but were consistently more abundant and frequent at the BBMO than in the subset of EukBank samples. These ASVs accounted for 17.8% of the community metabarcoding signal in BBMO (37.5% in May 2018 and 22.0% in September 2020) and 6.1% in EukBank. This suggests that these globally distributed species are better adapted to the Mediterranean coast.

**Fig. 3.**
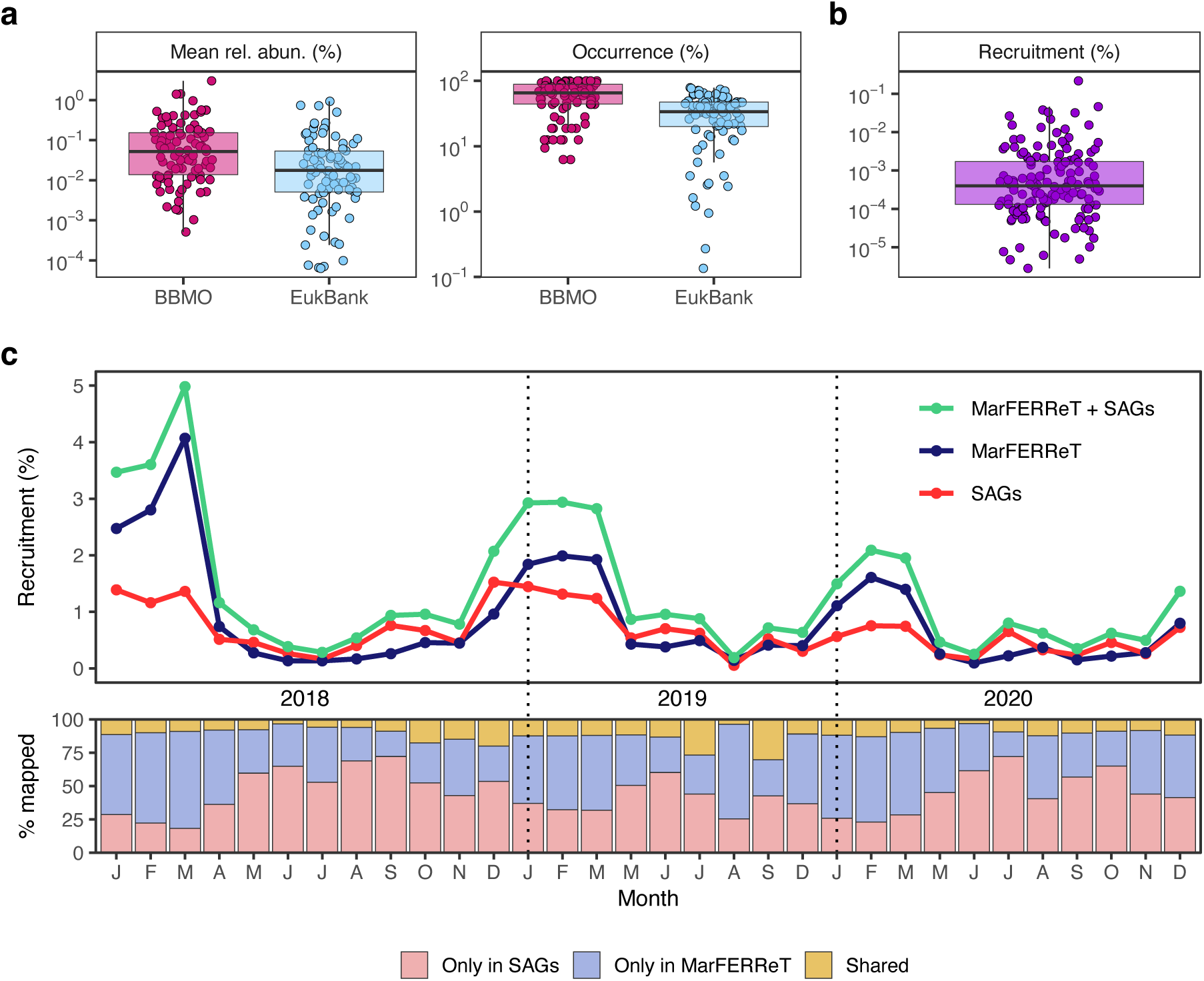
Ecological relevance of species with new single-cell genomes. **a**, Relative abundance and occurrence of ASVs corresponding to SAGs in marine picoplankton samples from the BBMO and EukBank metabarcoding datasets. **b,** Relative abundance of metagenomic reads recruited by each SAG across BBMO metagenomes (2018–2020). **c,** Added read recruitment in BBMO metagenomes by the SCG dataset and the MarFERRet protein catalogue. The lower panel shows the proportion of reads recruited exclusively by SAGs, exclusively by MarFERRet, or by both.

The second dataset used to evaluate the ecological relevance of the new genomes was a BBMO metagenomic time series of the picoplankton (0.2-3 µm cell fraction), dominated by prokaryotes and with a minor but significant protist contribution. All 147 predicted gene sets contributed to read recruitment, with a single SAG recruiting as much as 0.22% of the total (Fig. 3b). We also performed metagenomic read recruitment using MarFERReT, an extensive database of eukaryotic proteins^31^ (Fig. 3c). Across the three years, the two databases recruited similar proportions of reads (0.66% in SAGs and 0.86% in MarFERReT; 1.35% together), despite the stark difference in protein counts (1.6 million vs. 27.9 million proteins, respectively). Each dataset recruited distinct subsets of reads, with only a small fraction in common (Fig. 3c), highlighting a high complementarity between genomes obtained from single cells and from cultures. The largest contributor to the MarFERReT signal was *Bathycoccus prasinos* (first ASV in BBMO), while the largest contributor to the SAG signal was ICM0001 (second ASV in BBMO). The genome of ICM0001 was identical (ANI >99%) to that of the strain RCC434 recently deposited at NCBI. As RCC434 is the holotype for *Micromonas bravo*^32^, the strain CCAC1681 deposited in MarFERReT as *M. bravo* (ANI of 87.5% with RCC434) needs reclassification. *B. prasinos* and *M. bravo* drove the seasonal oscillations observed in metagenomic recruitment patterns (Fig. 3c). Overall, these results show that our new genomes provide strong resolving power for interpreting complex metagenomic datasets.

A selection of the most abundant BBMO species with SCG data based on metagenomic data is shown in Table S3. Photosynthetic taxa, primarily Mamiellophyceae but also Prymnesiophyceae, dominated the read recruitment, accounting together for 88% of the metaG signal. Among heterotrophs, MAST-4, MAST-1 and *Minorisa* were the most abundant. Five of these 20 dominant genomes appeared among the 20 most abundant ASVs, indicating clear concordance in these cases, but 15 dominant ASVs lacked a SCG representative (Table S3). Some of these absences were expected, as cells from groups like Ciliophora, Acantharea, and Cryptomonadales were not targeted during the sorting step. More striking was the high number of dominant MALV ASVs not recovered in our genomic dataset, consistent with reports of MALV overrepresentation in metabarcoding surveys^33^.

### Genomic architecture of the diverse marine protists

The analysis of dominant marine prokaryotes like *Pelagibacter* has revealed extensive genomic streamlining associated with large effective population sizes and adaptation to oligotrophic waters^34^, and this pattern has also been observed in the picoeukaryotic species *Micromonas*^35^. Here, we sought to determine how general this pattern is among the largely uncultured protist species represented by our newly generated genomes. To this end, we estimated the genome size based on assembly completeness and quantified the relative proportions of exonic, intronic, and intergenic regions. Benchmarking against three SAGs corresponding to cultured species showed strong agreement between our predicted genome sizes and those of reference strains (Table S4). Regarding the genomic regions, inferred from genomic and transcriptomic data in reference strains, our predicted values and reference annotations showed again high concordance, supporting the robustness of our SCG-based genome architecture estimates.

To explore patterns across taxa, we performed a principal component analysis (PCA) using genome size and the proportions of exonic, intronic, and intergenic regions (excluding two divergent genomes, a dinoflagellate and a ciliate). K-means clustering of the PCA space resolved five major genome-architecture types (Fig. 4a), each delineated by specific genomic features (Figs. 4b, 4c). Cluster 1 included small, compact genomes with very high exonic content (∼80%). Cluster 2 also displayed small genomes but with a higher proportion of intergenic regions (∼70% exonic and ∼25% intergenic). Clusters 3 and 4 exhibited intermediate genome sizes and lower exonic content (∼55–65%), with 3 enriched in intronic and 4 in intergenic regions. Finally, cluster 5 encompassed the largest genomes, characterized by low exonic content and high levels of both intronic and intergenic regions.

**Fig. 4.**
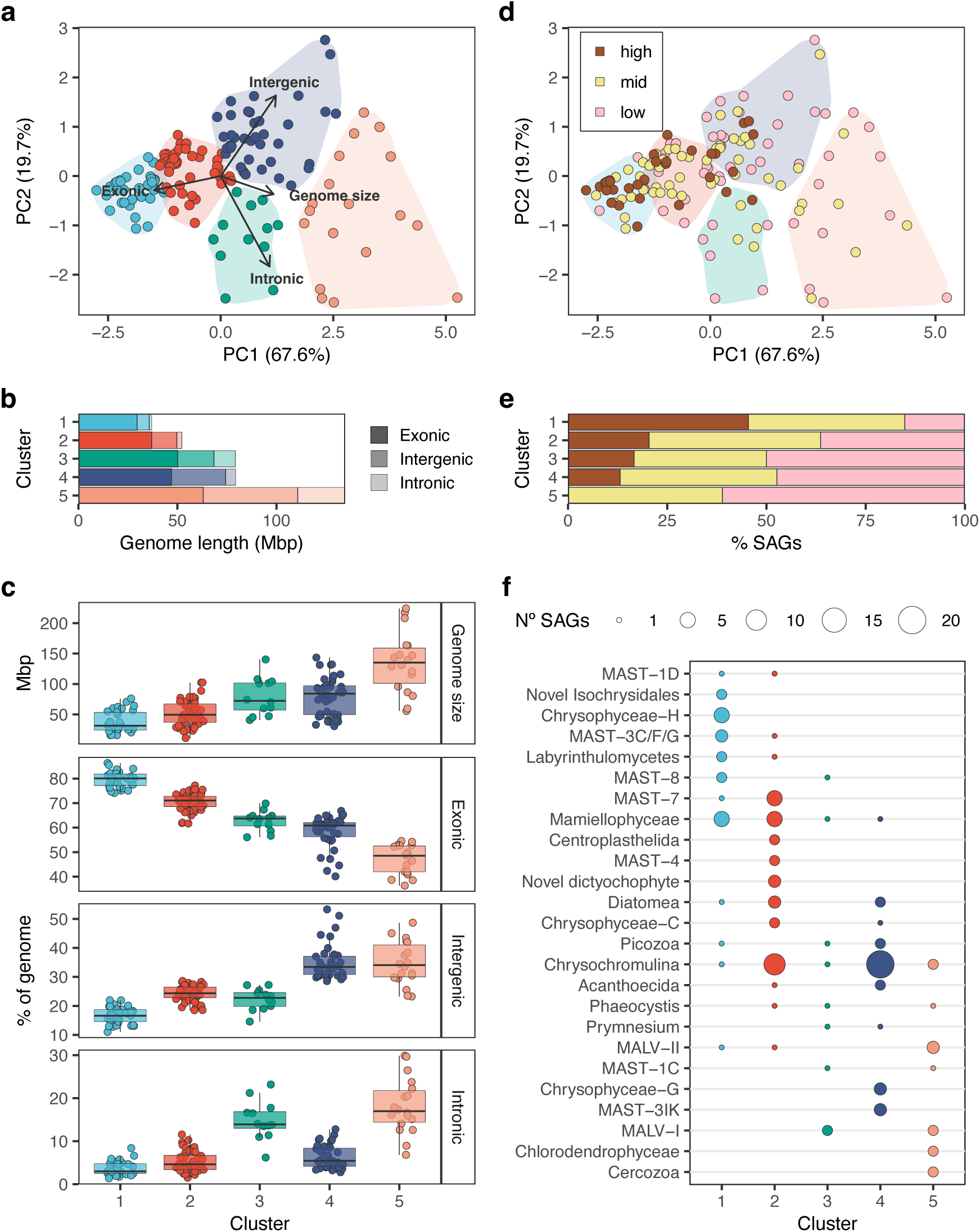
Genomic architecture of the final genomes. **a**, Principal component analysis (PCA) based on estimated genome size and proportions of exonic, intronic and intergenic regions, revealing five clusters. **b,** Mean genome architecture for the five clusters. **c,** Distribution of the four genomic features across the five clusters. **d,** Same PCA plot than panel **a** with points coloured by corrected relative abundance in BBMO metagenomes (high >0.005%; medium 0.001–0.005%; low <0.001%). **e,** Contribution of high, medium and low abundant genomes in each of the five clusters. **f,** Distribution of genomes of defined taxonomic groups among the 5 clusters.

Genome architecture was closely linked to ecological success. When mapping in the PCA plot the relative abundance of each species in the BBMO dataset, dominant species strongly accumulated in clusters 1 and 2 (Fig. 4d), which contained the majority of the 20 most abundant species (10 and 8, respectively; Table S3). The contribution of high abundant taxa gradually declined from cluster 1 that contained the highest proportion to cluster 5, which included none (Fig. 4e). Thus, the most abundant protist species in Blanes Bay typically possess small, compact genomes. Genomic streamlining, especially in intronic regions, seems to be a prerequisite for protist dominance in marine ecosystems. Nonetheless, genome compactness alone was not sufficient for high abundance, as many species within clusters 1 and 2 still remained rare.

Taxonomic distribution across clusters generally showed consistent pattern at lower ranks, with related species grouping together (Fig. S4) and accumulating in a given cluster (Fig. 4f). Cluster 1 contained all members of Chrysophyceae-H and the new Isochrysidales clade, and the majority of MAST-3 (C, F and G), MAST-8, and Labyrinthulomycetes. It also contained many Mamiellophyceae species. Cluster 2 included most *Micromonas* species (except the dominant one that was in cluster 1), most Diatomea, and all members of Centrohelida and a novel Dictyochophyceae clade. About half of the *Chrysochromulina* species belonged to cluster 2. Regarding heterotrophs, this cluster aggregated the abundant MAST-4 and most MAST-7 species. Cluster 3 contained the lowest number of species, with a heterogeneous representation of taxonomic groups. Cluster 4 was dominated by photosynthetic species with large genomes, including the remaining *Chrysochromulina* species and Chrysophyceae-G, together with the nano heterotrophs MAST-3 (K and I), Picozoa, and Acanthoecida choanoflagellates. Finally, cluster 5 included most MALVs, Cercozoa, Telonemia and Chlorodendrophyceae.

### Functional enrichment analysis and key gene families

About half of the predicted genes from SAGs had an annotated function, yielding a rich functional dataset for multiple analyses. As an initial assessment of the functional resolution of our partial genomes, we searched for functions enriched when comparing pigmented versus colourless cells, and vice versa (Table S5). All species coexist in the same habitat and share similar cell sizes, but they differ in their trophic strategy, with a clear dichotomy between chloroplast-containing cells and colourless cells. As expected, the genomes of pigmented cells were enriched in GO terms related to photosynthesis, biosynthetic processes, and ion and metabolite transport (Fig. 5a). Conversely, heterotrophic species were enriched in functions associated with cytoskeleton organization and motility, cell adhesion, signal transduction, and environmental sensing, reflecting an active lifestyle based on resource detection, prey capture and digestion.

**Fig. 5.**
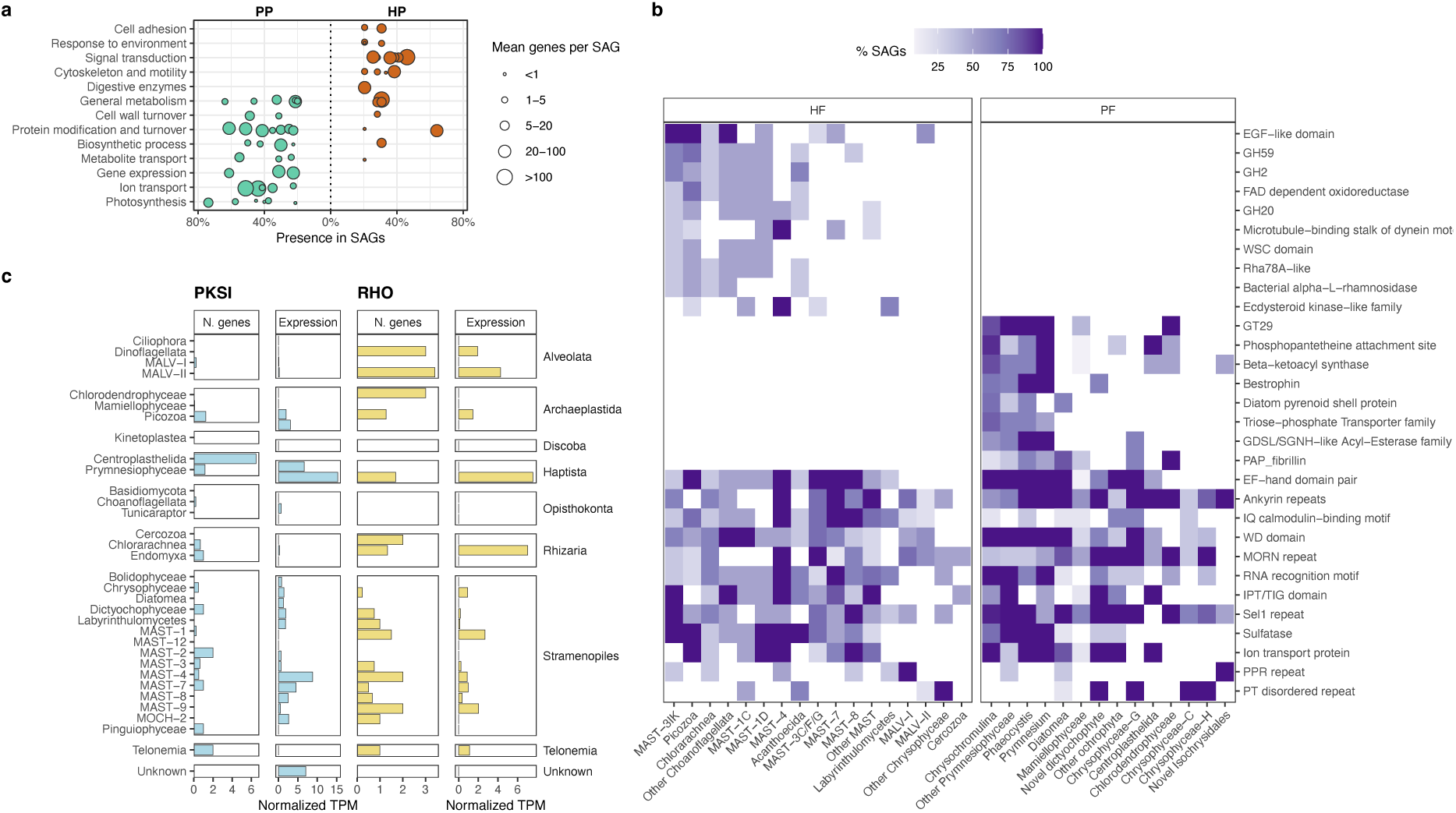
Functional analyses with the final genomes. **a**, Dots represent Gene Ontology Biological Process terms enriched in heterotrophic compared with phototrophic protists (HP; orange) and in the opposite comparison (PP; green). GO terms are grouped into broad functional categories; dot sizes indicate the number of genes per SAG. The list of enriched GO terms is shown in Table S5. **b,** Selection of Pfam domains enriched in SAGs as compared to the EukProt database of protist proteins. They are separated by being enriched in HP, in PP, or in all cells. Colour indicate the percentage of SAGs in each subgroup with the domain enriched. **c,** Number of polyketide synthase I (PKSI) and rhodopsin (RHO) genes per SAG within taxonomic groups. Expression levels of these genes in BBMO metatranscriptomes, normalized by metagenomic read abundance per taxonomic group.

In a second step, to identify functions that might be essential for protist success in the marine environment, we searched for Pfam domains enriched in our species compared with protist proteomes in EukProt. From the first list of 1313 enriched domains, we kept those enriched in several species from at least two taxonomic subgroups (Fig. S5). Some of these 303 domains were enriched in heterotrophic or photosynthetic taxa only, and others did not show a clear trend. Representative domains illustrating prevalent functional attributes are shown in Fig. 5b. Marine protists were mainly enriched in two functional categories: tandem-repeat proteins involved in protein binding and macromolecular complex assembly (Ank, Sel1, WD40), and domains involved in signalling and regulation like calcium signalling (EF-hand and IQ calmodulin) or kinase-mediated pathways. Other enriched functions were related to adhesion (IPT/TIG), RNA recognition (RRM), and cytoskeleton organization and motility (MORN). Regarding trophic strategies, heterotrophic taxa showed enrichment in the degradation of complex polysaccharides (Bac_rhamnosidase), prey/polysaccharide attachment (EGF-like and Rha78-like), motility and signalling. In contrast, photosynthetic species were enriched in glycan anabolism, plastid regulation, ion homeostasis, and an expanded repertoire of secondary metabolism pathways, likely involved in toxin production and antimicrobial defence.

Within this context, polyketide synthase (PKS) domains were particularly enriched in several photosynthetic groups, notably Prymnesiophyceae. PKS enzymes catalyse the synthesis of polyketides, a diverse class of secondary metabolites with antibiotic, antitumor, antifungal, and immunosuppressive activities^36^, also playing putative roles in microbial competition and fitness^37^. We assessed the genomic potential for polyketide production by identifying PKS genes (Table S6). Type I PKS genes, the most important of the three PKS families, were broadly distributed across taxa (Fig. 5c), with Centroplasthelida harbouring the highest number per SAG, followed by Telonemia, MAST-2, and Picozoa. Beyond genomic presence, PKS genes were actively expressed in BBMO metatranscriptomes (Fig. 5c), highlighting their functional relevance in situ.

We finally searched for genes encoding rhodopsins (RHO), a diverse family of light-responsive proteins involved in energy transduction or environmental sensing^38^ for which an increasing number of functional types are being described across the tree of life^39^. RHO genes were widespread in most groups but absent in Opisthokonta, Picozoa, MALV-I, Centroplasthelida, and Diatomea (Fig. 5c, Table S6). Phylogenetic placement within a reference framework showed that most of the genes clustered in three functional types (Fig. S6). Channel rhodopsins, functioning as light-gated ion channels with putative sensory roles, were more frequent in phototrophs and also occurred in MALV and MAST lineages. Proton-pump rhodopsins were found mainly in heterotrophic taxa (MALV-II, MASTs, and Telonemia) and also in Prymnesiophyceae (potentially mixotrophs), suggesting a role in phagosome acidification. Enzyme rhodopsins that activate downstream enzymatic responses were found in Prymnesiophyceae, Mamiellophyceae, and Chlorodendrophyceae. As with PKS genes, rhodopsins were transcriptionally active, pointing to previously unrecognized phototrophic or sensory capabilities in marine protists.

## Discussion

Single-cell genomics (SCG) in microbial ecology emerged around 20 years ago^12^ and is now an established component of the methodological toolkit for prokaryotes^40^. Comparable work on microbial eukaryotes followed shortly thereafter^16,41^ and despite proposed analytical pipelines^42^, the approach is still not widely used to generate *de novo* genomes of uncultivated species^43^. This limitation stems in part from the broad range of eukaryotic genome sizes, with some lineages possessing genomes of several gigabases^44^, for which SCG is expected to be challenging. Yet, SCG is promising for genomes up to 100-200 Mb, especially when MDA biases are mitigated by coassembling conspecific cells. In this work we applied SCG to capture dominant small-sized marine protists, representing one of the first efforts at this scale. Pigmented protists were single-cell sorted based on their pigments, while colourless protists were sorted after DNA staining^45^, which proved effective according to the consistent cytogram signatures of cells from the same species. Previous studies selected the cells to sequence by 18S rDNA screening^15,19^, but we show here that this step overlooks some taxa, thereby hindering genomic analysis of dominant members. Our data also suggest that avoiding cryopreservation may increase WGA recovery and genome completeness, while recent advances in WGA protocols further ensure more complete genome recovery^46^. Overall, the genomes obtained here encompass multiple eukaryotic lineages and together capture a large fraction of the diversity of the natural community.

This dataset reveals unique genomic trends linked to ecological success, in which streamlining is a recurrent mechanism suggesting convergent evolution across distinct taxonomic backgrounds. The most striking example of this pattern is found in distinct picophotosynthetic taxa with very reduced genomes. Mamiellophycean species (*Micromonas*, *Ostreococcus*, and *Bathycoccus*) have high GC-content, which entails greater nitrogen and phosphorus investment^47^. In contrast, Chrysophyceae-H and the new Isochrysidales group present low GC content, with the former enriched in phosphorous transporter domains and the latter having the PPR repeat associated with plastid control as one of the few enriched domains. This suggests that picophotosynthetic lineages have evolved distinct genomic strategies to cope with nutrient limitation. In particular, mamiellophycean species may tolerate higher nutrient demands due to their physiological efficiency, whereas Chrysophyceae-H and the new Isochrysidales appear better adapted to nutrient deficiency. Thus, these picoalgal groups differentially streamlined their gene content to evolve different functional adaptations. These patterns highlight that genome streamlining is not uniform but instead reflects lineage-specific trade-offs between nutrient economy, cellular function, and ecological specialization.

In marine heterotrophic organisms, genomic streamlining also occurs, with a tendency toward the reduction of non-coding regions but with a less pronounced decrease in genome size and gene content. This is likely because heterotrophs require a broader functional repertoire to sustain their living, including genes for motility, signalling, adhesion, and environmental sensing. We also detect the enrichment of protein domains associated with the degradation of complex polysaccharides across lineages (MASTs, Picozoa, choanoflagellates). This finding suggests that some phylogenetically distant heterotrophic lineages may have convergently evolved similar functional repertoires, facilitating their adaptation to apparently similar ecological niches.

Overall, our work provides an ecologically relevant genomic dataset of key protist players in marine ecosystems. Even with partial genomes, our SCG data enable robust characterization of genomic architecture and represent a critical advance toward resolving the genomic and functional diversity of dominant and often uncultured protists. Combined with accurate taxonomic classification and protein enrichment analyses, our data establish a robust framework for identifying both convergent and lineage-specific adaptations and for generating hypotheses about survival mechanisms in oligotrophic marine systems. Moreover, our new genomes can be linked to metagenomic and metatranscriptomic datasets, offering an appropriate taxonomic context for their interpretation. We have also expanded the range of taxa harbouring polyketide synthase and rhodopsin genes, opening new avenues to investigate their ecological roles in marine ecosystems. Finally, data recovered from poorly known lineages may contribute to studies of deep eukaryotic evolution, while the entire dataset may inform future cultivation efforts and single-cell interaction studies, including predation, symbioses, and viral infections.

## Methods

### Single cell sorting, whole genome amplification and Illumina sequencing

Sampling was conducted at the Blanes Bay Microbial Observatory (BBMO) in the NW Mediterranean. Surface seawater was filtered at sea through a 200 µm nylon mesh, transported to the lab, and single cells were sorted upon arrival (less than 4 hours after sampling) at the Flow Cytometry Unit of the CRG (Center for Genomic Regulation, Barcelona). Cell sorting was done in a BD Influx cell sorter (Becton & Dickinson Biosciences, San Jose, CA) that had been exhaustively cleaned for two days for DNA free assays^48^. A first sample (March 6, 2018) was used to optimize cell sorting gates. Phototrophic Protists (PP) were detected in an unstained aliquot based on chlorophyll (PECY5 channel, 670/30nm BP-filter) and phycobilin autofluorescence (PE channel, 580/40nm BP-filter). A second aliquot was stained with SYBR Green I (1:10000 initial stock) for 10 minutes to detect Heterotrophic Protists (HP) in the same channels: PE records the stain signal and PECY5 is roughly related to cell size^45^. HP and PP cells of different size were sorted as distinct subpopulations (pico/nano). To verify the performance of these gates (PP/HP and pico/nano), we sorted 300-1000 cells per gate into a tube with 1% glutaraldehyde. Cells were stained with DAPI, placed on a 0.2 µm pore-size polycarbonate filter and visualized by epifluorescence microscopy. Sorted cells were quantitatively recovered by microscopy, with sizes 1-3 µm in the pico and 3-6 µm in the nano fraction.

A first SCG effort was conducted with a sample collected on May 8, 2018. Single cells were placed into wells containing 2 µL of PBS buffer of a 96-well plate. The first column was left for negative controls (no cells) and the last for positive controls (5-10 cells). Sorted cells were indexed with their PE/PECY5 values. Four plates were prepared (1 for pico-PP [80 cells], 2 for pico-HP [160 cells] and 1 for nano-HP [86 cells]), sealed and kept at −80°C until processed. Whole genome amplification (WGA) from single cells in the four plates was done at the University of Alicante. First, cells were lysed for 10 minutes with Direct Lysis Buffer (Qiagen), followed by adding a Stop Solution (Qiagen). Then, WGA was done by multiple displacement amplification (MDA) with MDA-X phi 29 polymerase^24^ as described before^49^. Each well had SYTO9 stain to follow real time DNA amplification in a fluorescence plate reader (CLARIOStar, BMG LabTech). Reagents for cell lysis and MDA were treated to avoid amplifying contaminant DNA^48^. Reactions were stopped when many cells from the plate reached a maximum, typically after 3-4 hours. Positive WGA occurred in 244 out of the 326 cells. These 244 single amplified genomes (SAGs) were subjected to a PCR with the 18S rDNA eukaryotic primers 528F and Euk-B^25^ and PCR products were Sanger sequenced with the forward primer. Illumina libraries with KAPA HyperPrep (Roche) were prepared for 24 SAGs per sorted population. Nextera libraries were also prepared for one third of the SAGs; both libraries performed similarly in terms of assembly size and completeness (data not shown). Illumina sequencing was done in a HiSeq 2500 platform (2×250 PE) at the CNAG (Centre Nacional d’Anàlisi Genòmica) with a sequencing depth of ∼1.6 Gb per SAG. Based on the taxonomic classification given by the 18S rDNA, 9 additional SAGs were sequenced in a MiSeq platform (2 x 250 PE), totalling 81 cells sequenced in May 2018.

After the positive performance of the first test, we conducted a second single cell sorting effort with a sample collected on September 15, 2020. On that day, 17 plates were prepared as before (4 for PP [pico+nano], 8 for pico-HP and 5 for nano-HP). Cells sorted as PP were treated separately as pico-PP and nano-PP based on their cytogram indices (Fig. S7). Some plates were sent to UoA for WGA, and 384 cells were chosen for Nextera libraries and sequencing in a NovaSeq 6000 platform (2×150 PE) at the CRG. Some plates were sent to the Genomics Core Leuven (https://www.genomicscore.be/) for whole genome amplification with PicoPLEX (Takara), and 372 cells were chosen for Nextera libraries and sequencing in a NovaSeq platform (2×150 PE).

### SAG assembly, species delineation and co-assembly

Illumina reads from the 837 cells sequenced were subjected to the following process. Raw reads were trimmed with TrimGalore v0.6.10 (github.com/FelixKrueger/TrimGalore) with default settings and a minimum length of 75 bp. Genome assembly was performed using SPAdes v4.2.0^50^ with the options --sc and -k 21,33,55,77,99,127. Contigs <1 kb were discarded from the initial assembly. Completeness was estimated with BUSCO v5.3.2^51^, which searched for a set of 255 conserved single-copy eukaryotic genes from OrthoDB v11. Contigs >3 kb were assigned to bacteria, archaea, or eukaryotes (nuclear or organelle) by Tiara v1.0.3^52^. Based on these statistics, we selected the SAGs to be kept for downstream analysis. For the 2018 batch, 68 of the 81 SAGs gave acceptable genomes (4 were bacteria and 9 were too small). The 2020-UoA batch (384 cells), yielded a substantial number of bacteria in the HP plates, and we kept 227 SAGs. The 2020-Leuven batch (372 cells) was largely contaminated, and we only kept 30 SAGs. Gene predictions in the final set of 325 SAGs were generated with BRAKER v.3.0.7^53^ using OrthoDB version 11 as protein evidence. General taxonomy of the predicted proteins was assigned with eggNOG-mapper v2.1.10^54^ and Kaiju v1.10.1^55^, and functional annotation was done with InterProScan v5.76-107.0^56^. Genomes were decontaminated by removing contigs >3 kb originating from prokaryotes, bacteria or archaea based on Tiara results, and by removing contigs 1-3 kb in size that derive from prokaryotic genomes based on the gene annotations.

For taxonomic identification of the 325 resulting SAGs and to determine if some were from the same species, three steps were applied. First, the 18S rDNA sequence was retrieved by a blastn search of the assembly against the PR2 database v5.1.1^57^. Second, the proteins of each SAG were searched using blastp against the proteins from all SAGs, which allowed identifying groups of SAGs sharing genes. Third, these groups of SAGs were subjected to average nucleotide identity (ANI) comparisons with a custom script, which fragments each subject genome into 1 kb segments before BLAST. SAGs belonging to the same species (virtually identical 18S rDNA and very high ANI) were co-assembled with SPAdes. Coassembled genomes (COSAGs) were decontaminated as before. The final curated genomes from SAGs and COSAGs were subjected to a final gene prediction with BRAKER, and were functionally annotated with eggNOG-mapper and InterProScan. Final BUSCO analyses were done with predicted genes.

### Genome mapping against metabarcoding and metagenomic datasets

Environmental DNA at the BBMO was collected monthly from January 2018 to December 2020 (32 samples). About 10 litres of 200 µm-prefiltered surface seawater were filtered using a peristaltic pump through a 20 µm nylon mesh and polycarbonate filters of 3 µm and 0.2 µm pore size. Filters of the pico-size fraction (0.2-3 µm) were kept at −80°C until DNA extraction^22^. Metabarcoding analysis targeted the V4 region of the 18S rDNA amplified with eukaryotic primers^58^. PCR products were sequenced at Allgenetics on a MiSeq platform (2×250 PE). Raw reads were processed with DADA2^59^ to delineate ASVs (Amplicon Sequence Variants). ASVs were classified to broad taxonomic groups using the eukaryotesV4 database^60^, and ASVs from metazoa, streptophyta and nucleomorphs were removed before further analyses. Additional metabarcoding datasets analysed here included the Ocean Sampling Day (OSD), which targeted protists < 200 µm in coastal marine samples^29^, and the EukBank, the largest compendium of natural samples from diverse habitats^30^. Already-processed OSD data were retrieved from metaPR^2^ tool^61^.

In these metabarcoding datasets, we searched for ASV that were identical to the rDNA of our single cell derived genomes. This could be done in 109 out of the 147 genomes, as 14 lacked the 18S rDNA, and 24 had this gene but not the V4 region. This resulted in 104 identical matches in the BBMO dataset. There were 7 pairs of genomes with the same ASV (4 Prymnesiophyceae, 1 MALV-II, 1 Mamiellophyceae and 1 Picozoa). In some cases, these pairs differed outside the V4 region, but in other cases they had identical complete 18S rDNA, so they stand as examples of the limited taxonomic resolution of the 18S rDNA gene. The ANI among these pairs was about 85% (first 7 plots in Fig. S1b), confirming that they were different genomes with identical 18S rDNA. Nevertheless, the fact that in most cases each genome had a unique ASV supports the power of the 18S rDNA (and the V4 region) to delineate species.

Metagenomes were obtained by sequencing the same DNA extracts from the BBMO time series in an Illumina NovaSeq6000 (2×150 PE) at the CNAG, with a sequencing depth of 35 Gb per sample. The taxonomic composition of the metagenomes was assessed with the pipeline previously described^60^, by which Illumina reads encompassing the V4 region of the 18S rRNA gene were first extracted through a permissive blastn and then classified to an eukaryotic group or supergroup. For metagenomic read recruitment, reads of 135 bp or larger were kept and mapped with DIAMOND v2.0.7^62^ against the collection of proteins from SAGs and the MarFERReT v1.1.1 database (after removing 7 SAGs). Alignments longer than 45 amino acids with 100% identity were retained. For reads with more than one hit, a consensus taxonomy was computed using a simple last common ancestor algorithm that removed conflicting taxonomic levels. The number of recruited reads was normalized by the total number of reads in each metagenomic sample. To quantify the expression of SAG genes, Salmon v1.10.2^63^ was used in mapping-based mode to align 21 BBMO metatranscriptomes collected in 2021–2022 against SAG nucleotide coding sequences. A table of transcripts per million (TPM) values was obtained.

### Phylogenomics, genomic architecture and functional exploration of the SCG data

Phylogenomic reconstructions were conducted with PhyloFisher version 1.2.14 using the dataset of 240 marker proteins derived from 304 eukaryotic species representing the full breadth of eukaryotic diversity^64^. The obtained super-matrix was used to generate a first maximum-likelihood (ML) tree with IQ-TREE version 3.0.1^65^ with the model LG+C20+F+G, which was then used as a guide for the final ML tree, performed with the model LG+C60+F+G+PMSF and 1000 ultrafast bootstrap, and 1000 SH-aLRT replicates.

We used a custom script to extract different genomic regions (exonic, intronic and intergenic) using the BRAKER gene annotations from the final assemblies. In addition, the estimated genome size of each species was calculated in base of a putative 100% BUSCO completeness. To test if these predictions from SAGs were reliable, we compared the values obtained in three SAGs belonging to protist species whose genomes had been fully sequenced and annotated with RNA-seq data: *Bathycoccus prasinos*, *Phaeocystis cordata* and *Ostreoccoccus* sp. RCC809 (Table S4). This showed that ICM0077 was virtually complete.

GO term enrichment analyses were based on InterProScan annotations and conducted with package topGO version 2.62.0^66^ in R version 4.2.2, by comparing the genomes of pigmented species against those of unpigmented ones. This was done by conducting an enrichment analysis of each genome against the opposing group, and then identifying the genes and functions that were consistently enriched in species from the same trophic mode. The functional enrichment was performed on SAGs with at least 15% completeness, applying the following threshold: occurrence in at least 20% of the cells within the trophic group, and enrichment factor of 3 (three times more enriched in one group relative to the other).

Pfam domain enrichment analysis was done with a custom script by comparing the protein domains of each of the 147 genomes against a background file summarizing the weighted average protein domain composition of the EukProt protist proteomes^67^ (excluding animals, plants and marine SAGs). A domain was considered enriched in SAGs when passed the significance threshold in a Fisher exact test (p-value <0.05). To look for commonly enriched protein domains across the dataset, we selected for those enriched in at least 5% of the species from any of the PP and HP fractions and found in at least two different taxonomic subgroups.

Polyketide synthases (PKS) and rhodopsins (RHO) genes within the SAGs were searched based on Pfam annotations. Three types of PKS genes were inspected. Type I PKS genes were considered complete when they contained three functional domains: ACP, acyl carrier protein (PF00550, PF14573, or PF23297); AT, acyl transferase (PF00698); and KS, ketoacyl synthase (PF00109, PF02801, or PF16197). They were considered incomplete when they displayed the KS domain but lacked 1-2 of the other domains. For Type II PKS, condensation domain (PF00668) was searched. For Type III PKS, chalcone and stilbene synthase domains (PF00195, PF02797) were searched. For RHO, the bacteriorhodopsin-like protein domain (PF01036) was searched.

## Data availability

Raw Illumina sequencing data from SAGs are available at the European Nucleotide Archive (ENA) under accession PRJEB108838. The final dataset of 147 genomes is available at Zenodo (https://doi.org/10.5281/zenodo.18786764), which display a folder per SAG with the assembly, genome analyses, gene predictions and gene annotations. Raw Illumina amplicon data from the BBMO time series are available at ENA under accession PRJEB97769. Raw Illumina metagenomic data from the BBMO series are available at ENA under accession PRJEB51979. Metatranscriptomic data used here is available at Zenodo (https://doi.org/10.5281/zenodo.18922885). Data to reproduce the analyses and figures are available at GitHub (https://github.com/MassanaLab/sags_bbmo_paper).

## Code availability

All code used to build the SAG dataset and reproduce the analyses and figures is available at GitHub (https://github.com/MassanaLab/sags_bbmo_paper).

## Acknowledgements

This work was supported by the Spanish projects DIVAS (PID2019-108457RB-I00) and EPIC (PID2022-137508NB-I00), the “Severo Ochoa Centre of Excellence” accreditation (CEX2019-000928-S), the project BYGENEX (ERC-2024-AdG, EU, 101198091) and the EASI-Genomics TransNational Access project MAPSAG (PID10388 Grant Agreement No 824110). We thank the Genomics Core Leuven facility, in particular Kristine Stepanyan and Wim Meert. We are grateful to Alexandre Bote Tronchoni and Mario Gómez Álvarez for their contribution in lab work. Bioinformatic analyses were carried out in Marbits (ICM-CSIC; https://marbits.icm.csic.es/) and FinisTerrae III (CESGA; https://www.cesga.es/) computer platforms.

## References

1. Bar-On, Y. M. & Milo, R. The biomass distribution on Earth. Proc. Natl. Acad. Sci. USA 115, 6506–6511 (2019).

2. Caron, D. A., Countway, P. D., Jones, A. C., Kim, D. Y. & Schnetzer, A. Marine protistan diversity. Annu. Rev. Mar. Sci. 4, 467–493 (2012).

3. Worden, A. Z., Follows, M. J., Giovannoni, S. J., Wilken, S., Zimmerman, A. E. & Keeling, P. J. Rethinking the marine carbon cycle: Factoring in the multifarious lifestyles of microbes. Science 347, 1257594 (2015).

4. Massana, R. Eukaryotic picoplankton in surface oceans. Annu. Rev. Microbiol. 65, 91–110 (2011).

5. Massana, R. et al. Phylogenetic and ecological analysis of novel marine stramenopiles. Appl. Environ. Microbiol. 70, 3528–3534 (2004).

6. Guillou, L. et al. Widespread occurrence and genetic diversity of marine parasitoids belonging to Syndiniales (Alveolata). Environ. Microbiol. 10, 397–408 (2018).

7. Not, F. et al. Picobiliphytes: A marine picoplanktonic algal group with unknown affinities to other eukaryotes. Science 315, 252–254 (2007).

8. Caron, D. A. et al. Probing the evolution, ecology and physiology of marine protists using transcriptomics. Nat. Rev. Microb. 15, 6–20 (2017).

9. Hoegh-Guldberg, O., Northrop, E., & Lubchenco, J. The ocean is key to achieving climate and societal goals. Science 365, 1372–1374 (2019).

10. Nishimura, Y. & Yoshizawa, S. The OceanDNA MAG catalog contains over 50,000 prokaryotic genomes originated from various marine environments. Sci. Data 17, 305 (2022).

11. Delmont, T. O. et al. Functional repertoire convergence of distantly related eukaryotic plankton lineages abundant in the sunlit ocean. Cell Genomics 2, 100123 (2022).

12. Stepanauskas, R. & Sieracki, M. E. Matching phylogeny and metabolism in the uncultured marine bacteria, one cell at a time. Proc. Natl. Acad. Sci. USA 104, 9052–9057 (2007).

13. Mangot, J. F. et al. Accessing to the genomic information of unculturable oceanic picoeukaryotes by combining multiple single cells. Sci. Rep. 7, 41498 (2017).

14. Strassert, J. F. H. et al. Single cell genomics of uncultured marine alveolates shows paraphyly of basal dinoflagellates. ISME J. 12, 304–308 (2018).

15. Schön, M. E. et al. Single cell genomics reveals plastid-lacking Picozoa are close relatives of red algae. Nat. Commun. 12, 6651 (2021).

16. Martínez-García, M. et al. Unveiling in situ interactions between marine protists and bacteria using single cell genomics. ISME J. 6, 703–708 (2012).

17. Wittmers, F. et al. Symbionts of predatory protists are widespread in the oceans and related to animal pathogens. Cell Host & Microbe 33, 182–199 (2025).

18. Wideman, J. G. et al. Unexpected mitochondrial genome diversity revealed by targeted single-cell genomics of heterotrophic flagellated protists. Nat. Microbiol. 5, 154–165 (2020).

19. Labarre, A. et al. Comparative genomics reveals new functional insights in uncultured MAST species. ISME J. 15, 1767–1781 (2021).

20. López-Escardó, D. et al. Reconstruction of protein domain evolution using single-cell amplified genomes of uncultured choanoflagellates sheds light on the origin of animals. Philos. Trans. R. Soc. B Biol. Sci. 374, 20190088 (2019).

21. Chen, W. et al. The hidden genomic diversity of ciliated protists revealed by single-cell genome sequencing. BMC Biol. 19, 264 (2021).

22. Giner, C. R. et al. Quantifying long-term recurrence in planktonic microbial eukaryotes. Mol. Ecol. 28, 923–935 (2019).

23. Giovannoni, S. J., Thrash, J. C. & Temperton, B. Implications of streamlining theory for microbial ecology. ISME J. 8, 1553–1565 (2014).

24. Stepanauskas, R. et al. Improved genome recovery and integrated cell-size analyses of individual uncultured microbial cells and viral particles. Nat. Comm. 8, 84 (2017).

25. Sieracki, M. E. et al. Single cell genomics yields a wide diversity of small planktonic protists across major ocean ecosystems. Sci. Rep. 9, 6025 (2019).

26. Pierella Karlusich, J. J., et al. A robust approach to estimate relative phytoplankton cell abundances from metagenomes. Mol. Ecol. Resour. 23, 16–40 (2023).

27. del Campo, J., Not, F., Forn, I., Sieracki, M. E., Massana, R. Taming the smallest predators of the oceans. ISME J. 7, 351–358 (2013)

28. Ruiz-Trillo, I., Kin, K. & Casacuberta, E. The origin of metazoan multicellularity: A potential microbial black swan Eevent. Annu. Rev. Microbiol. 77, 499–516 (2023).

29. Kopf, A. et al. The ocean sampling day consortium. GigaScience 4, s13742–015–0066–5 (2015).

30. Berney, C., Mahé, F., Henry, N., Lara, E. & de Vargas, C. EukBank 18S V4 dataset. Zenodo (2023).

31. Groussman, R. D., Blaskowski, S., Coesel, S. N. & Armbrust, E. V. MarFERReT, an open-source, version-controlled reference library of marine microbial eukaryote functional genes. Sci. Data 10, 926 (2023).

32. Simon, N. et al. Revision of the genus *Micromonas* Manton et Parke (Chlorophyta, Mamiellophyceae), of the type apecies *M. pusilla* (Butcher) Manton & Parke and of the species *M. commoda* van Baren, Bachy and Worden and description of two new Sspecies based on the genetic and phenotypic characterization of cultured isolates. Protist 168, 612–635 (2017).

33. Massana, R. et al. Marine protist diversity in European coastal waters and sediments as revealed by high-throughput sequencing. Environ. Microbiol. 17, 4035–4049 (2015).

34. Giovannoni, S. J. et al. Genome streamlining in a cosmopolitan oceanic bacterium. Science 309, 1242–1245 (2005).

35. Worden, A. Z. et al. Green evolution and dynamic adaptations revealed by genomes of the marine picoeukaryotes *Micromonas*. Science 324, 268–272 (2009).

36. Jenke-Kodama, H., Sandmann, A., Müller, R., Dittmann, E. Evolutionary implications of bacterial polyketide synthases. Mol. Biol. Evol. 22, 2027–2039 (2005).

37. Kohli, G. S., John, U., Van Dolah, F. M. & Murray, S.A. Evolutionary distinctiveness of fatty acid and polyketide synthesis in eukaryotes. ISME J. 10, 1877–1890 (2016).

38. Pinhassi, J., DeLong, E. F., Béjà, O., González, J. M. & Pedrós-Alió, C. Marine bacterial and archaeal ion-pumping rhodopsins: Genetic diversity, physiology, and ecology. Microb. Mol. Biol. Rev. 80, 929–954 (2016).

39. Galindo, L.J., et al. Apusomonad rhodopsins: A new family of ultraviolet to blue light–absorbing rhodopsin channels. Proc. Natl. Acad. Sci. USA 122, e2510619122 (2025).

40. Pachiadaki, M. G. et al. Charting the complexity of the marine microbiome through Single-Cell Genomics. Cell 179, 1623–1635 (2019).

41. Yoon, H. S. et al. Single-cell genomics reveals organismal interactions in uncultivated marine protists. Science 322, 714–717 (2011).

42. Ciobanu, D. et al. A single-cell genomics pipeline for environmental microbial eukaryotes. iScience 24, 102290 (2021)

43. Schoenle, A. et al. Protist genomics: Key to understanding eukaryotic evolution. Trends Genet. 42, 869–882. (2025).

44. Lin, S. A decade of dinoflagellate genomics illuminating an enigmatic eukaryote cell. BMC Genomics 25, 932 (2024).

45. Zubkov, M. V., Burkill, P. H. & Topping, J. N. Flow cytometric enumeration of DNA-stained oceanic planktonic protists. J. Plankton Res. 29, 79–86 (2007).

46. Bowers, R. M. et al. scMicrobe PTA: near complete genomes from single bacterial cells. ISME Comm. 4, ycae085 (2024).

47. Elser, J., Acquisti, C. & Kumar, S. Stoichiogenomics: the evolutionary ecology of macromolecular elemental composition. Trends Ecol. Evol. 26, 38–44 (2010)

48. Rinke, C. et al. Obtaining genomes from uncultivated environmental microorganisms using FACS-based single-cell genomics. Nat. Protoc. 9, 1038–48 (2014).

49. Martinez-Hernandez, F. et al. Single-virus genomics reveals hidden cosmopolitan and abundant viruses. Nat. Comm. 8, 15892 (2017).

50. Prjibelski, A., Antipov, D., Meleshko, D., Lapidus, A. & Korobeynikov, A. Using SPAdes De Novo Assembler. Curr. Protoc. Bioinformatics 70, e102 (2020).

51. Tegenfeldt, F. et al. OrthoDB and BUSCO update: annotation of orthologs with wider sampling of genomes, Nucleic Acids Res. 53, D516–D522 (2025).

52. Karlicki, M., Antonowicz, S. & Karnkowska, A. Tiara: deep learning-based classification system for eukaryotic sequences. Bioinformatics 38, 344–350 (2022).

53. Gabriel, L. et al. BRAKER3: Fully automated genome annotation using RNA-seq and protein evidence with GeneMark-ETP, AUGUSTUS, and TSEBRA. Genome Res. 34, 769–777 (2024).

54. Cantalapiedra, C. P., Hernández-Plaza, A., Letunic, I., Bork, P. & Huerta-Cepas, J. eggNOG-mapper v2: Functional Annotation, Orthology Assignments, and Domain Prediction at the Metagenomic Scale. Mol. Biol. Evol. 38, 5825–5829 (2021).

55. Menzel, P. et al. Fast and sensitive taxonomic classification for metagenomics with Kaiju. Nat. Comm. 7, 11257 (2016).

56. Jones, P. et al. InterProScan 5: genome-scale protein function classification, Bioinformatics 30, 1236–1240 (2014).

57. Guillou, L. et al. The Protist Ribosomal Reference database (PR2): a catalog of unicellular eukaryote Small Sub-Unit rRNA sequences with curated taxonomy. Nucleic Acids Res. 41, D597–D604 (2013)

58. Balzano, S., Abs, E. & Leterme, S. Protist diversity along a salinity gradient in a coastal lagoon. Aquat. Microb. Ecol. 74, 263–277 (2015).

59. Callahan, B. J. et al. DADA2: High-resolution sample inference from Illumina amplicon data. Nat. Methods 13, 581–583 (2016).

60. Obiol, A. et al. A metagenomic assessment of microbial eukaryotic diversity in the global ocean. Mol. Ecol. Resour. 20, 718–731 (2020).

61. Vaulot, D. et al. metaPR2: A database of eukaryotic 18S rRNA metabarcodes with an emphasis on protists. *Mol*. Ecol. Res. 22, 3188–3201 (2022).

62. Buchfink, B., Reuter, K. & Drost, H.-G. Sensitive protein alignments at tree-of-life scale using DIAMOND. Nature Meth. 18, 366–368 (2021).

63. Patro, R., Duggal, G., Love, M. I., Irizarry, R. A. & Kingsford, C. Salmon provides fast and bias-aware quantification of transcript expression. Nature Meth. 14, 417–419 (2017).

64. Tice, A. K. et al. PhyloFisher: A phylogenomic package for resolving eukaryotic relationships. PLOS Biol. 19, e3001365 (2021).

65. Wong, T. K. F., et al. IQ-TREE 3: Phylogenomic Inference Software using Complex Evolutionary Models. https://ecoevorxiv.org/repository/view/8916 (2025).

66. Alexa, A. & Rahnenführer, J. topGO: Enrichment Analysis for Gene Ontology. doi:10.18129/B9.bioc.topGO (2025).

67. Richter, D. J., et al. EukProt: A database of genome-scale predicted proteins across the diversity of eukaryotes. Peer Com. 2, e56 (2022).

