## Supplementary Figures for "Single-cell sequencing reveals genome streamlining and functional diversity in ecologically dominant marine protists"

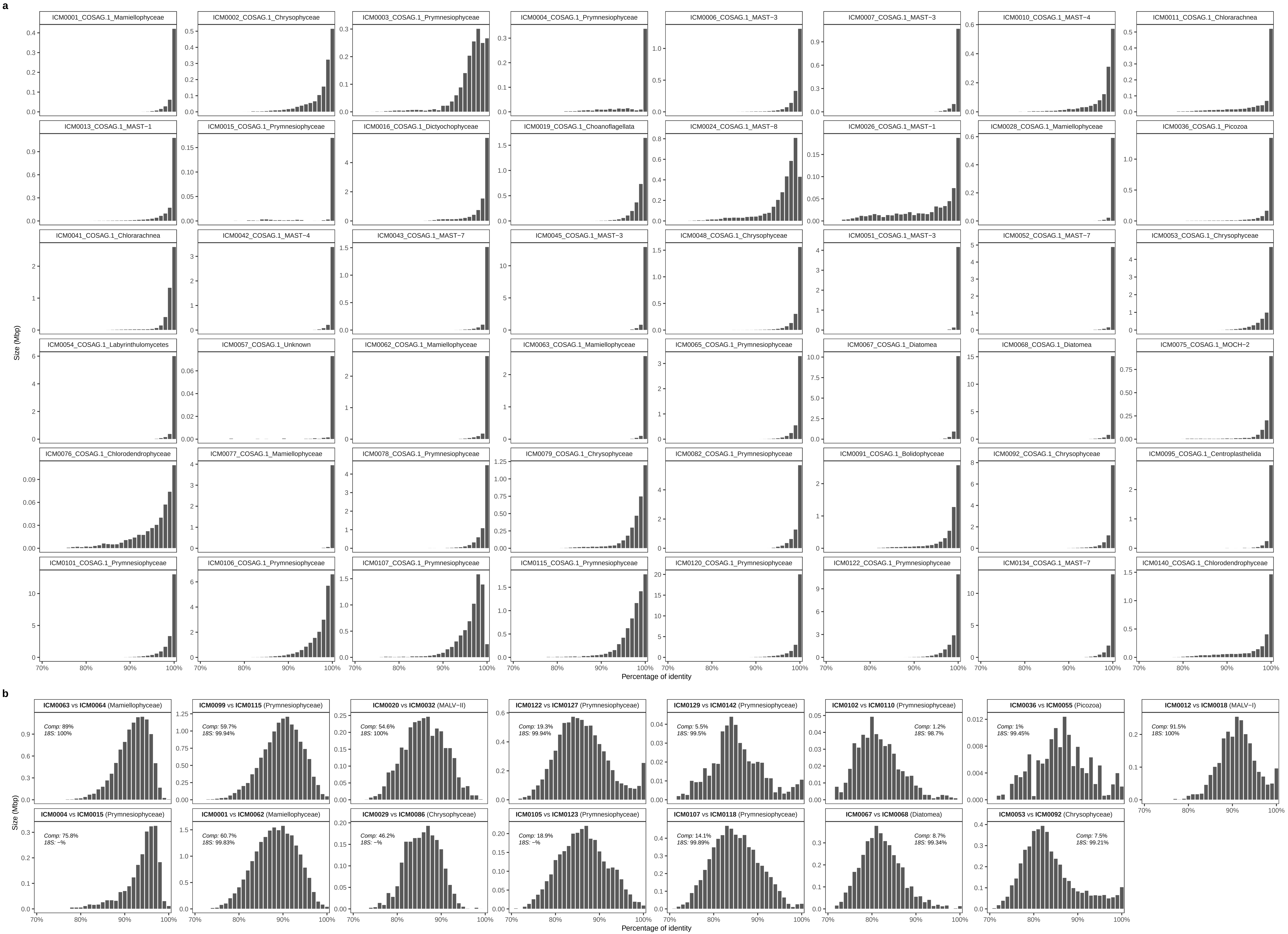


**Fig. S1 | Genome similarity among SAGs within each co−assembly (a) and between closely distinct genomes (b).** Pair−wise average nucleotide identity (ANI) comparisons were calculated by fragmenting the subject genome into 1 kb segments and using BLAST hits with a query >=0.5 kb. The similarity of the queried genome size was distributed in 1% bins histograms. Panels from (b), comparing different genomes, also display the sequence similarity of the 18S rDNA genes (the first 7 cases are those matching the same ASV) and the fraction of the genome that has been compared.

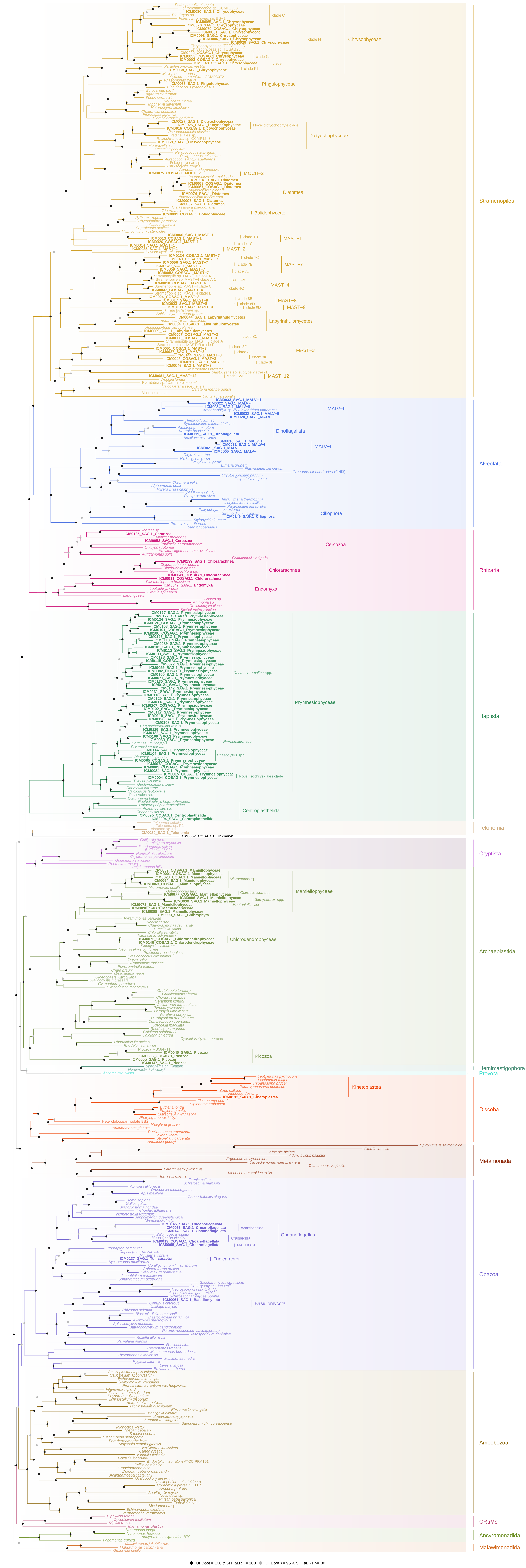


**Fig. S2 | Full multigene phylogenetic tree of the 147 final genomes and 304 reference genomes.** The tree was estimated in IQ−TREE with model LG+C60+F+G+PMSF, using a supermatrix of 240 conserved marker proteins constructed with PhyloFisher. SCG and reference genomes are coloured by supergroup. Taxonomic groups containing SCG are indicated. Bootstrap support is shown for all clades, with nodes highlighted in black when support is 100% and in grey when only slightly lower.

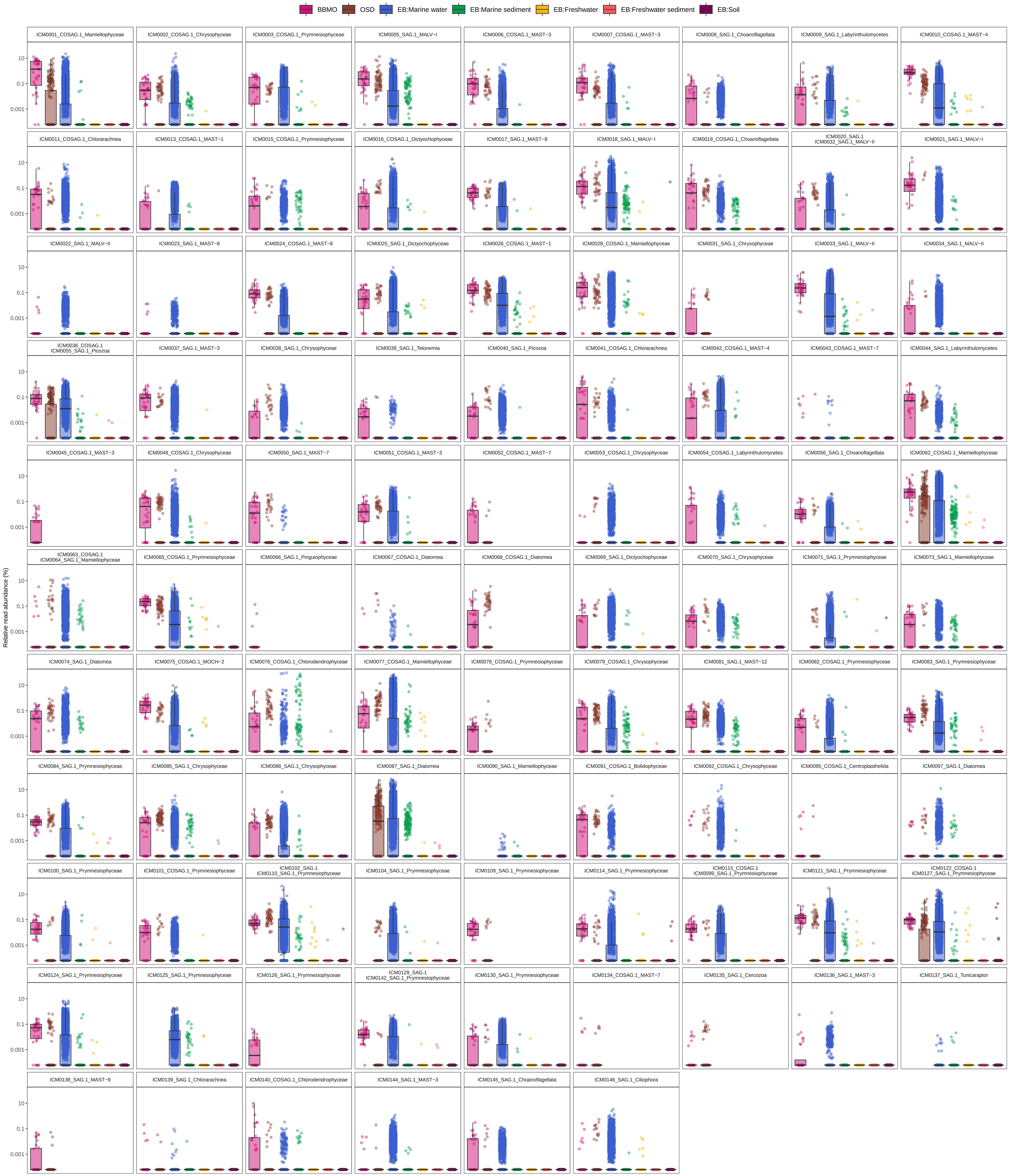


**Fig. S3 | Biogeography and habitat coverage of ASVs assigned to the final genomes.** Distribution of the relative abundance per sample of ASVs corresponding to the final genomes across large−scale metabarcoding datasets: BBMO (32 samples), OSD (319 samples), and EukBank subdivided into marine water (6545 samples), marine sediment (634 samples), freshwater (237 samples), freshwater sediment (153 samples) and soils (866 samples).

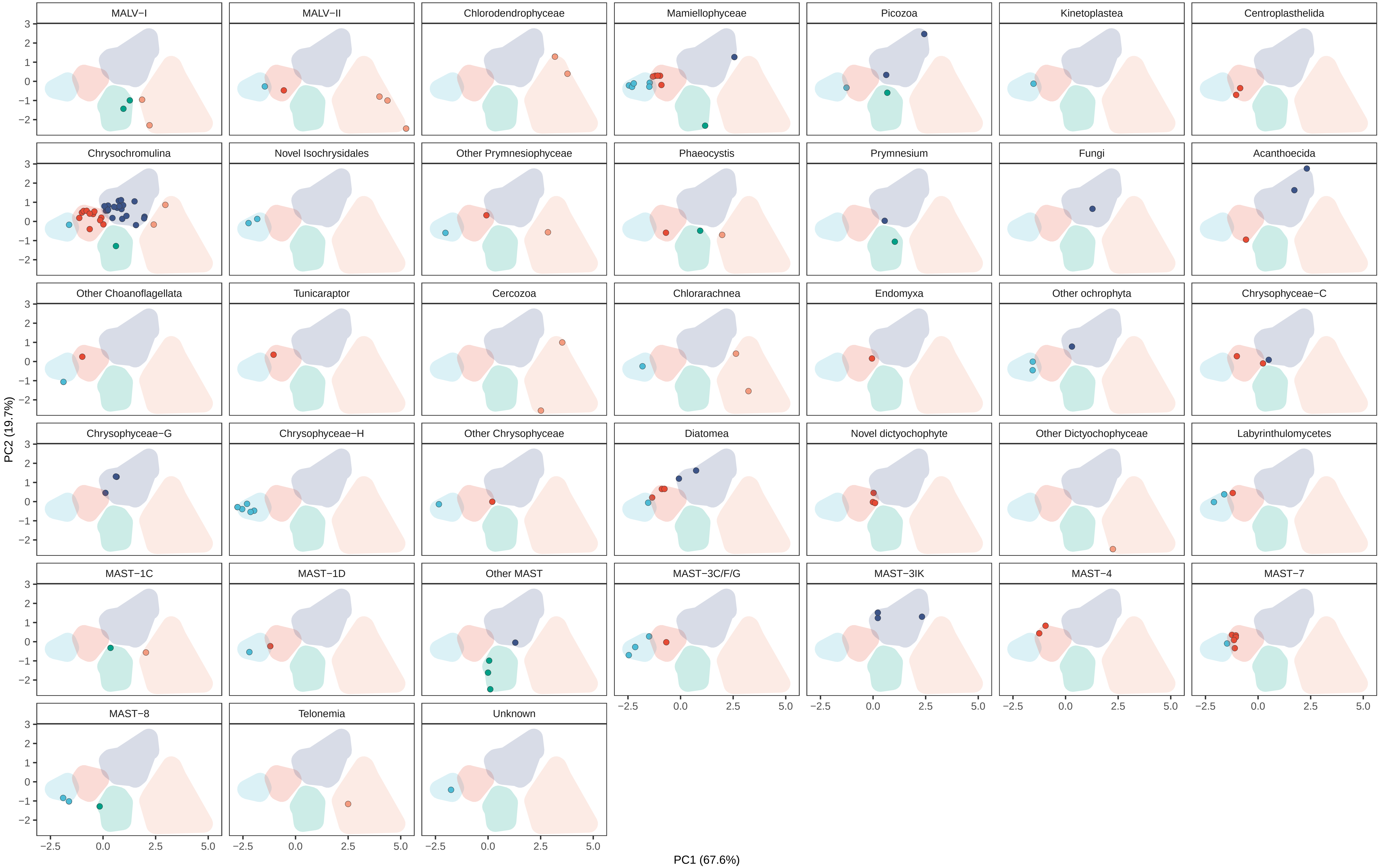


**Fig. S4 | Placement of the final genomes in the Fig. 4a PCA plot per taxonomic group.** Separate plots are presented for each distinct taxonomic group. Shaded areas indicate k−means clusters.

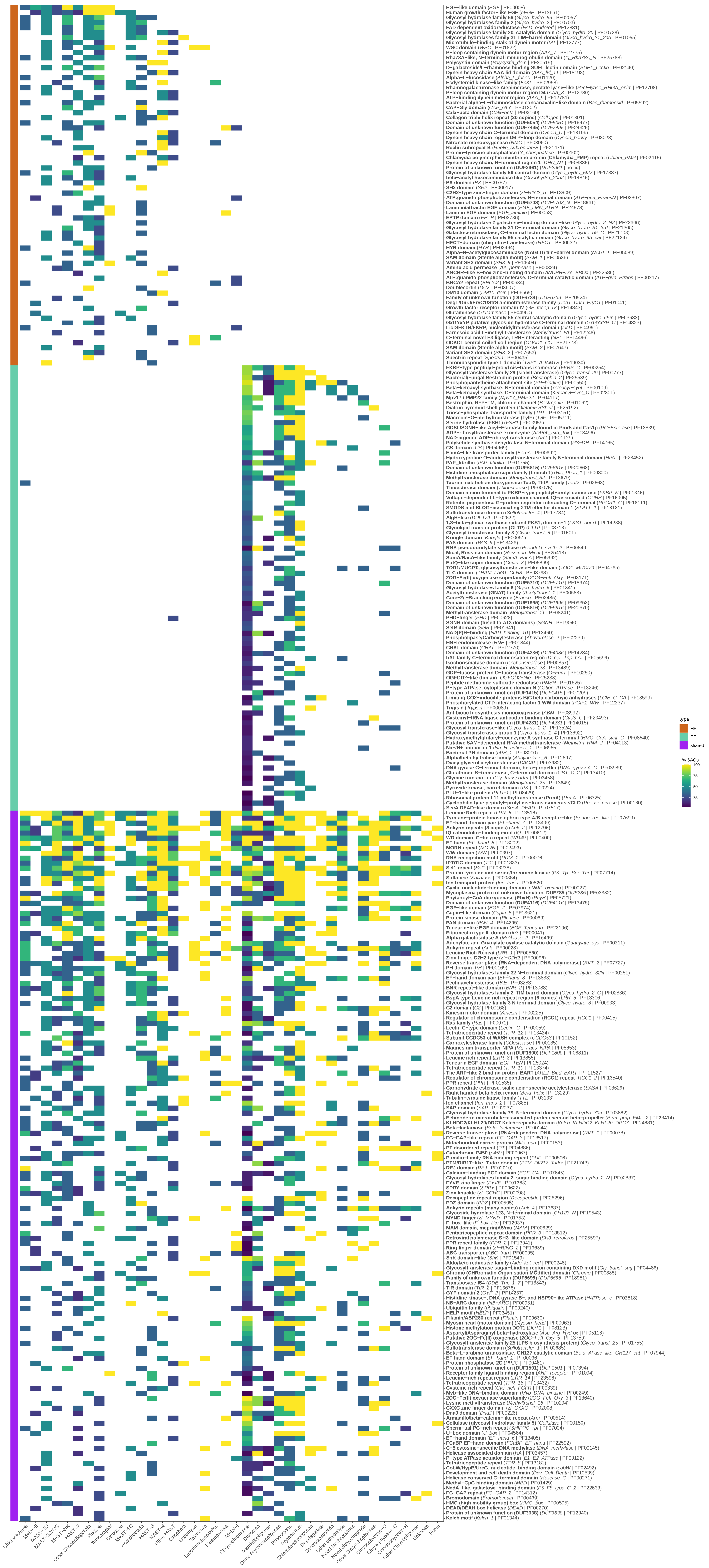


**Fig. S5 | Pfam domains enriched in final genomes as compared as the EukProt database of protist proteins.** The 303 selected domains are separated by being enriched only in HP, only in PP, or in both sets. Color intensity indicate the percentage of SAGs with the domain enriched in each subgroup.

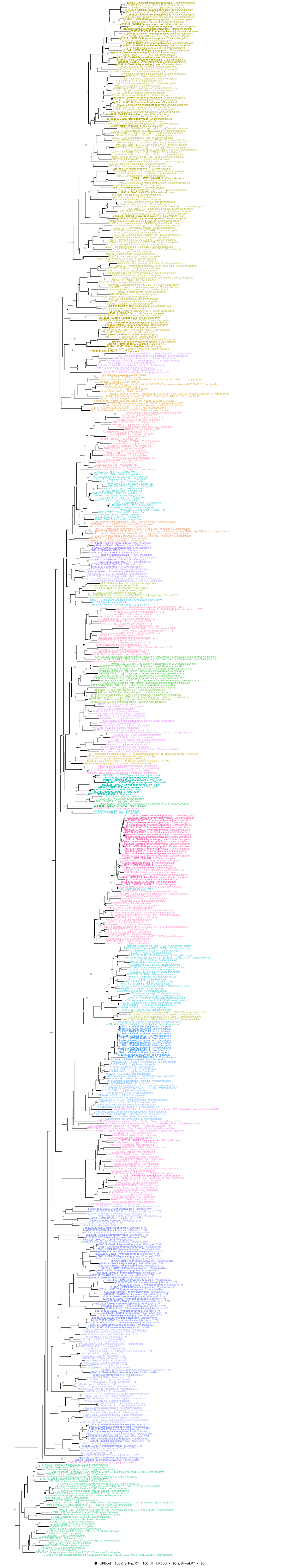


**Fig. S6 | Phylogenetic tree to place SCG rhodopsin genes.** Rhodopsin genes from SAGs were retrieved based on the Pfam domain PF01036 in gene annotations (158 genes; bold in the tree). For a phylogenetic tree they were combined with the sequences (and group delineation) presented in Galindo et al. 2025 and with 52 MarFERReT sequences (< 97% identity to the previous references). The corresponding group of each sequence is indicated after the name.

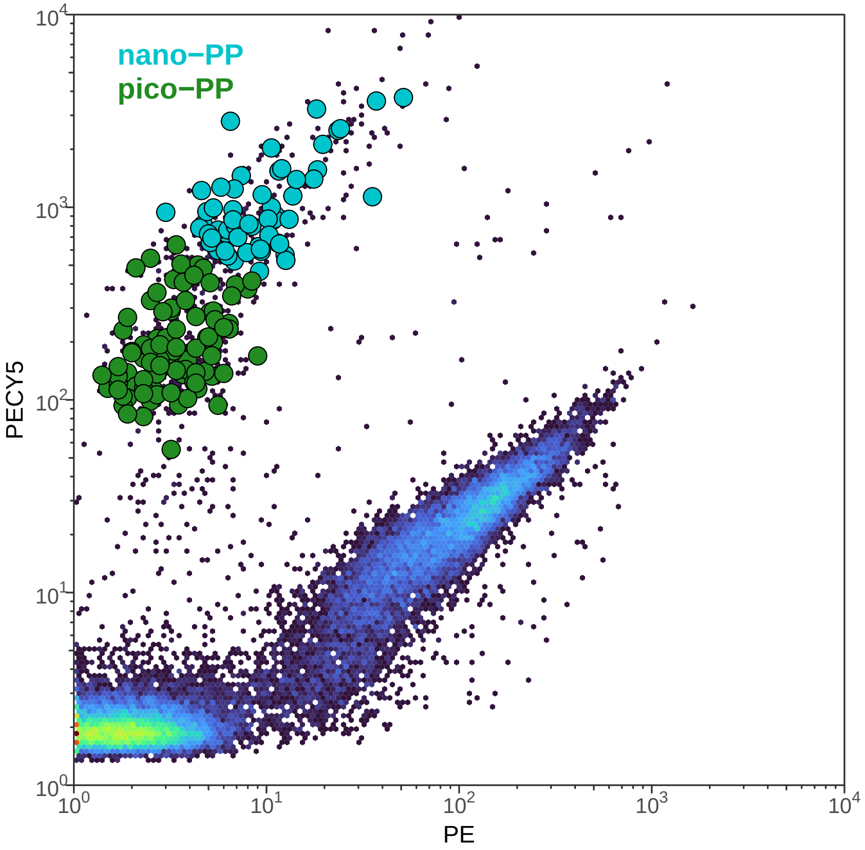


**Fig. S7 | Flow cytogram of the unstained sample collected in September 2020.** Large dots indicate sorted PP cells, which were subsequently classified as pico−PP and nano−PP based on index sorts.
